# Practical Design and Implementation of Animal Movements Tracking System for Neuroscience Trials

**DOI:** 10.1101/2020.07.26.221754

**Authors:** Majid Memarian Sorkhabi

## Abstract

**Background:** The nervous system functions of an animal are predominantly reflected in the behaviour and the movement, therefore the movement-related data and measuring behavior quantitatively are crucial for behavioural analyses. The animal movement is traditionally recorded, and human observers follow the animal behaviours; if they recognize a certain behaviour pattern, they will note it manually, which may suffer from observer fatigue or drift.

**Objective:** Automating behavioural observations with computer-vision algorithms are becoming essential equipment to the brain function characterization in neuroscience trials. In this study, the proposed tracking module is eligible to measure the locomotor behaviour (such as speed, distance, turning) over longer time periods that the operator is unable to precisely evaluate. For this aim, a novel animal cage is designed and implemented to track the animal movement. The frames received from the camera are analyzed by the 2D bior 3.7 Wavelet transform and SURF feature points.

**Results:** Implemented video tracking device can report the location, duration, speed, frequency and latency of each behavior of an animal. Validation tests were conducted on the auditory stimulation trial and the magnetic stimulation treatment of hemi-Parkinsonian rats.

**Conclusion/ Significance:** The proposed toolkit can provide qualitative and quantitative data on animal behaviour in an automated fashion, and precisely summarize an animal’s movement at an arbitrary time and allows operators to analyse movement patterns without requiring to check full records for every experiment.

## 1. INTRODUCTION

The function of the central nervous system has a direct effect on the motor behavior of animals and humans. Given that motor behavior is a qualitative parameter, it is important to convert this parameter into quantitative and measurable values [1]. If the accuracy of this measurement is acceptable, it will provide valuable data on the underlying neural circuit computation [2]. On the other hand, image/video processing and machine vision research have created a new branch in the biological sciences, called biological tracking or bio-tracking, which has vital applications in the fields of neuroscience, animal behavior, cognition and genetics sciences [1]. For example, to follow the effect of a drug on a laboratory mouse, its movements are monitored before and after treatment [3]. Most of the research utilized visual inspection by a human, which has low accuracy and may suffer from observer fatigue or drift [4].

Machine vision algorithms enable the operators to understand the meaning and content of images/videos by combining methods of image processing and machine learning tools [5]. Object tracking is an essential part of many applications of high-level machine vision, such as motion detection, brain-machine interface [6], video content indexing [7], traffic monitoring [8], and vehicle navigation, which are hot topics in research today.

The use of animal models allows researchers to do more research on specific disease conditions that are usually not possible in human experiments. However, researchers are trying to reduce the numbers of animal and do as small damage to the animals as possible [9].

Object tracking can be used for applications where the movements of objects are completely specific to the same type of object, and we can distinguish the identity and function of objects by the type of movement [10]. For example, identifying a person based on how they walk, or in medical applications for analysing movement disorders. For another instance, elderly people’s movement in a home can be explored, so that if a person falls on the ground for any reason, his/her condition is diagnosed by the object tracking system, and their condition can be reported to medical centres [11]. This modality is also utilized in sports applications to analyse the movement of players on the field [12].

In this research, 2D discrete Wavelet transform, and SURF feature points have been used to track animal movements. For this purpose, a special cage was built, and the required codes have been written in the MATLAB software. The details of the Wavelet transform can be found in [8]. SURF feature points are described below.

## 2. Techniques and methods

### 2.1 SURF feature points

The SURF feature points analyse the texture and pixel places to detect the maximum light reflection or the pixel changes, relative to the background, are remarkable. This method is used to extract high-level feature points. This algorithm has been used in the field of machine vision and watermarking [13]. What is important in this selection is the stability of the key points extracted by the proposed algorithm, so that it can correctly identify and model a specific feature of all images given to the algorithm. This method can also be implemented on the FPGA platform [14].

SURF points are generally shown as circles; the radius of each circle indicates the value of the SURF strength of that point, called descriptor; the larger the point radius, the stronger its SURF feature. A rule of thumb, the strength of each point of the SURF can be assumed to be the difference between the grey level of the pixel and that of the neighbouring pixels; the edges of the image, that have the maximum difference with their neighbouring pixels, have stronger SURF points. To test the SURF algorithm, we apply this method to two images with high and low interchanges, as illustrated in Figure 1. It can be seen that firstly this algorithm can find edge areas, secondly, the number of SURF feature points in images with higher edges is more. Therefore, using high-level SURF feature points can be a significant help in finding the desired points.

**Figure 1.**
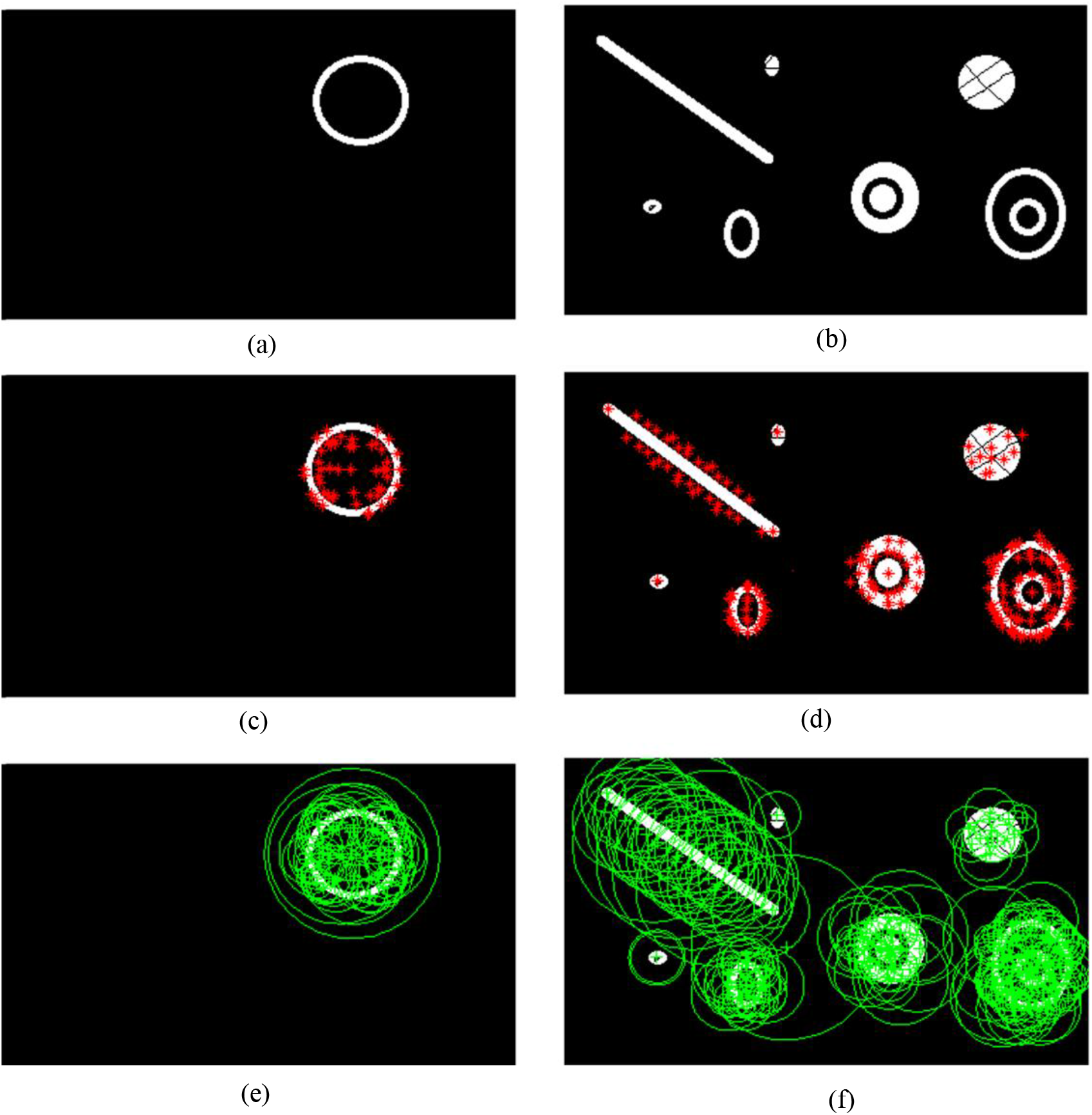
Using two test images for the SURF algorithm. (a) Test image #1 with low changes (edges). (b) Test image #2 with high changes (edges). (c) Location of SURF feature points for #1. (d) Location of SURF feature points for #2. (e) The size of SURF points (descriptor) for #1. (f) The descriptors for #2.

### 2.2 custom-made cage

The cage has an important effect on the behavior of the laboratory animal. Behaviours such as fear are directly related to the environmental conditions, including the temperature inside the cage, light, available food and water. Therefore, precise standards have been set for the construction of the cage, which must be considered in order to minimize the impact of the cage status on the behavior of the animal [15]. In this research, the size of 45 * 45 * 55 cm has been selected for the cage dimension, which can be seen in Figure 2. To control the animal’s environmental and movement conditions, the necessary hardware components were installed in the cage to monitor and adjust the temperature inside the cage, lighting and air circulation.

**Figure 2.**
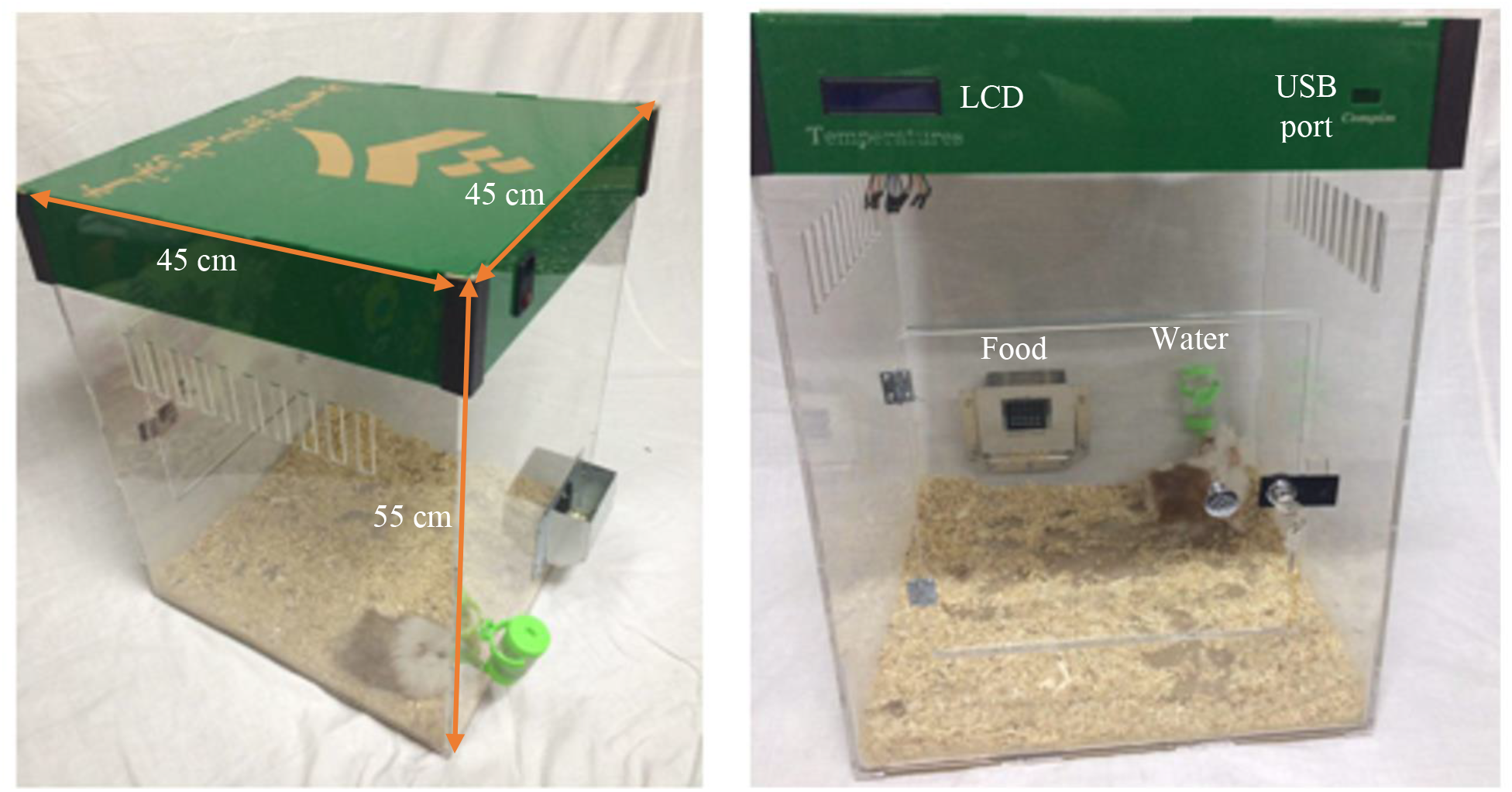
As-built cage in this research

The components of the cage are shown in Figure 3, including:

- Temperature sensor (LM35) located at the bottom of the cage.
- Heating element with a fan to control the temperature and the air circulation, which is located at the top of the cage.
- Lighting elements consisting of eight 1 W LEDs and a digital RGB camera.
- A speaker to create the required auditory stimulation in the relevant experiments.
- Electronic board with three relays, to control the lighting elements, and the 300 W heating element.
- An AVR microcontroller and an LCD, to control and display the parameters. This board connects to a computer with a TTL to USB converter.

**Figure 3.**
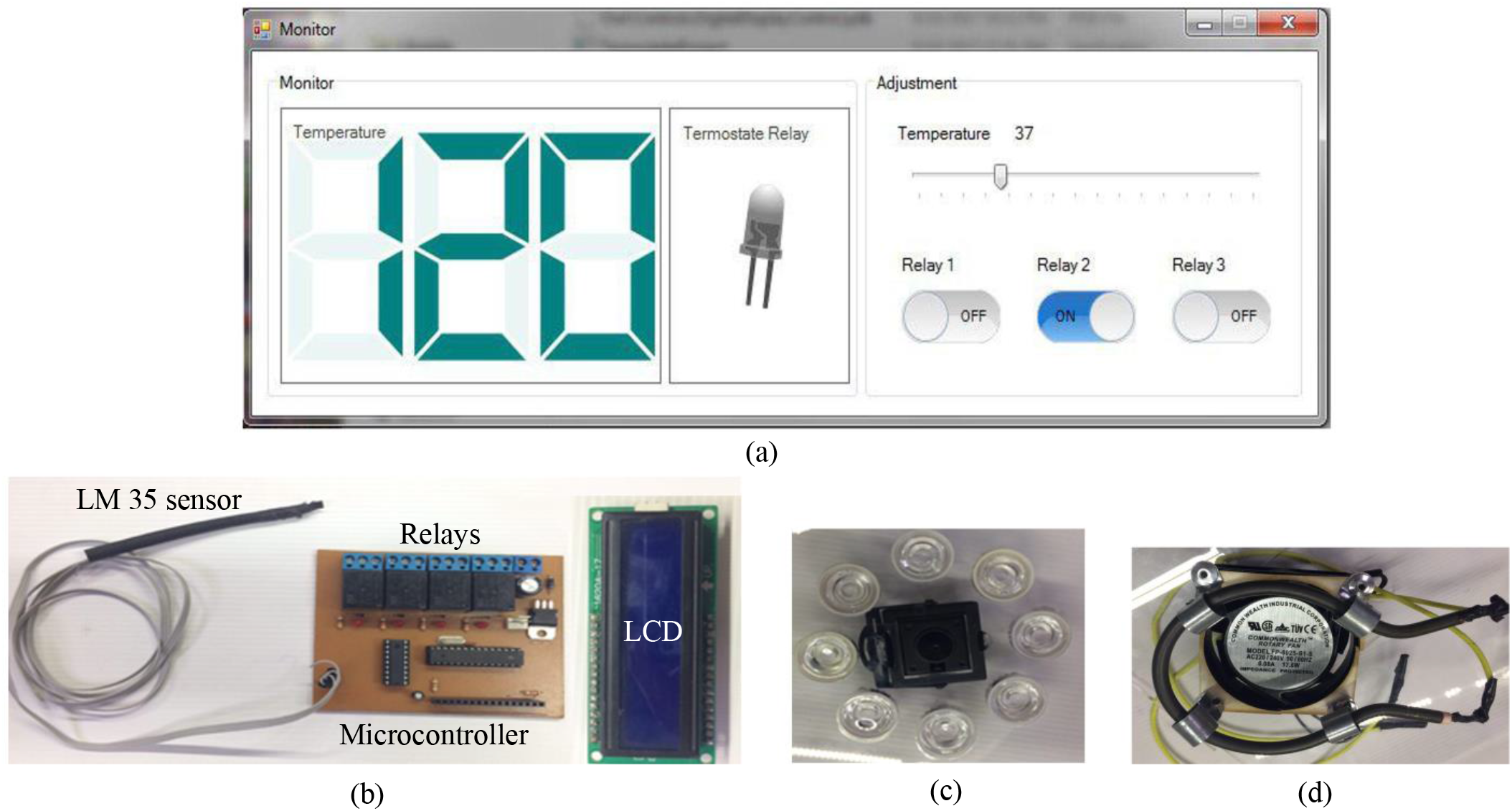
Cage hardware and user interface. (a) The user interface to monitor and control the lighting and the heating. (b) The temperature sensor, the AVR microcontroller, relays and the LCD. (c) LEDs and the RGB digital camera. (d) Heating element and the air circulation fan.

Using a custom-made program and the software interface, we can monitor the parameters inside the cage and change them if necessary. Fuzzy logic is used to control the temperature of the cage. Therefore, the environmental parameters of this cage are controlled completely automatically, and their accurate monitoring is possible.

### 2.3 Video processing and animal tracking

The video received from the camera inside the cage was used to study the movement behavior of the animal. By processing the obtained video, the necessary parameters such as animal’s movement path, eating/drinking time, location, duration, speed, frequency and latency of each behavior of an animal are extracted. To verify the research, a guinea pig (6 months old and male) was placed in a cage and its movements were analysed. Given that the color of the animal is white-brown and inhomogeneous, choosing this animal will better challenge the efficiency of the proposed algorithm and system. The video frames are RGB and can be captured at 25 frames per second.

The video processing steps are performed to search for the animal’s movement according to the block diagram in Figure 4. In the first step, all received frames are converted from RGB to the grey level. A frame of an empty cage (without an animal) is then taken and selected as the base frame. After placing the animal in the cage, the mainframes are received and subtracted from the base frame. This frame is called a differential frame. Then its two-dimensional discrete Wavelet transform is calculated. The approximate coefficients of this conversion are selected for use in the next step. Finally, the SURF feature points are calculated and the strongest point that has the most change concerning the whole frame is selected as the animal location.

**Figure 4.**
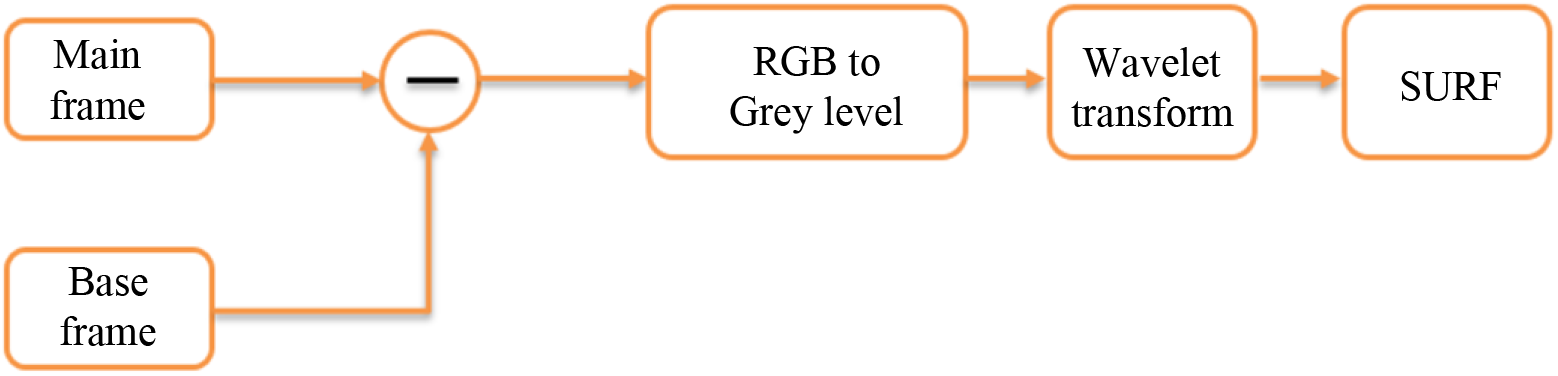
Video processing block diagram to extract the position of an animal

## 3- Results

### 3-1 Animal movement tracking

The base frame, the mainframe and the result of subtracting the base frame from the mainframe (differential frame) are shown in Figure 5. The value of the RGB to grey level conversion threshold is set to remove the extra image components. According to Figure 5, after the conversion, both the animal itself and the reflection of the animal remain on the wall of the cage.

**Figure 5.**
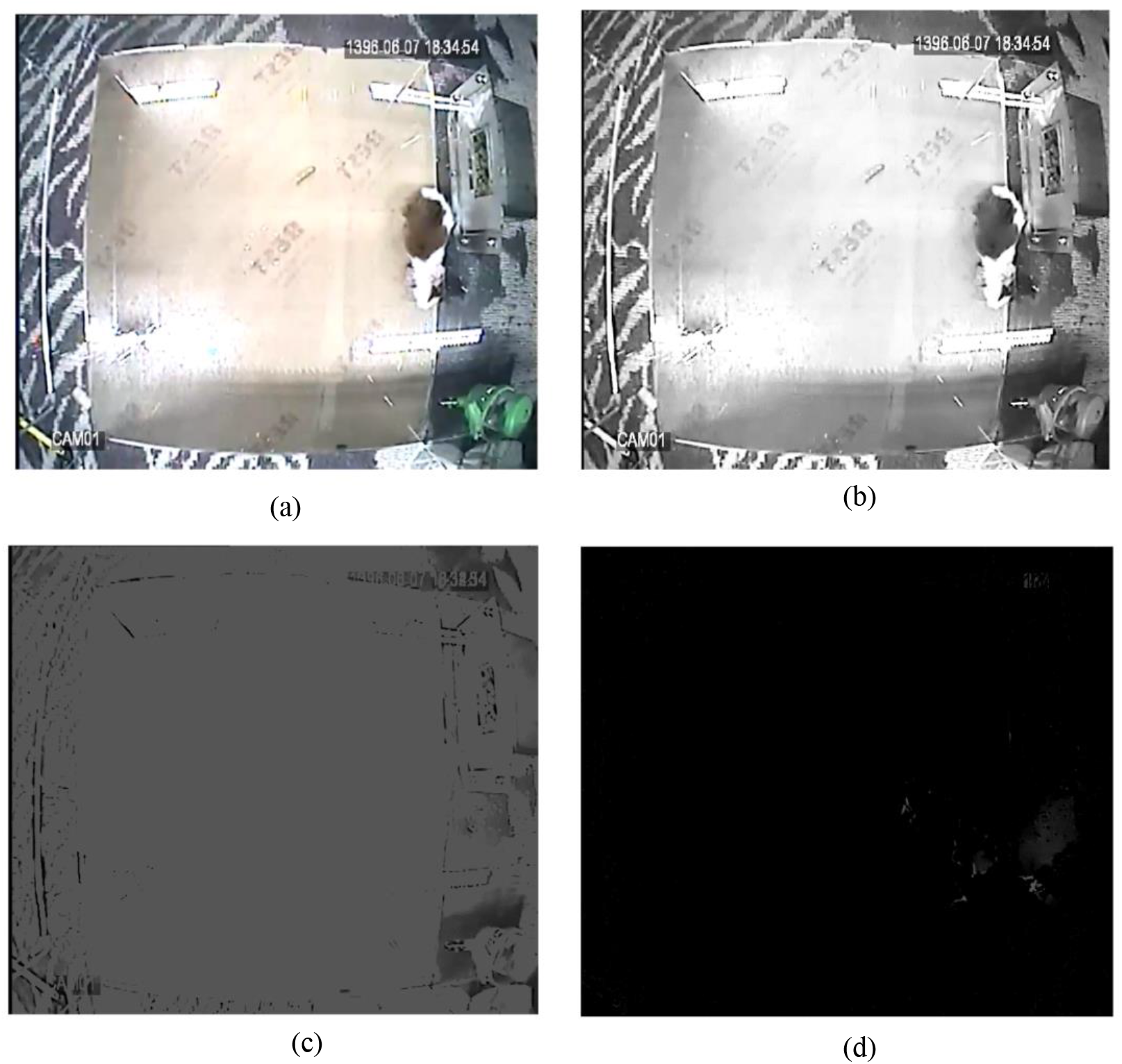
(a) Received RGB image (mainframe). (b) Convert RGB image to a grey level. (c) Base frame in the grey level. (d) The result of subtracting the base frame from the mainframe (differential frame)

The result of discrete Wavelet transforms and SURFT feature points of the differential frame are shown in Figure 6. Evidently, the strongest feature of the SURF is found on the wall of the cage (Figure 6(b)). To solve this problem, by completing the written code, we limit the selection of the SURF point to the cage floor. This only evaluates the floor points of the cage and shows the position of the animal inside the cage (Figure 6(c)).

**Figure 6.**
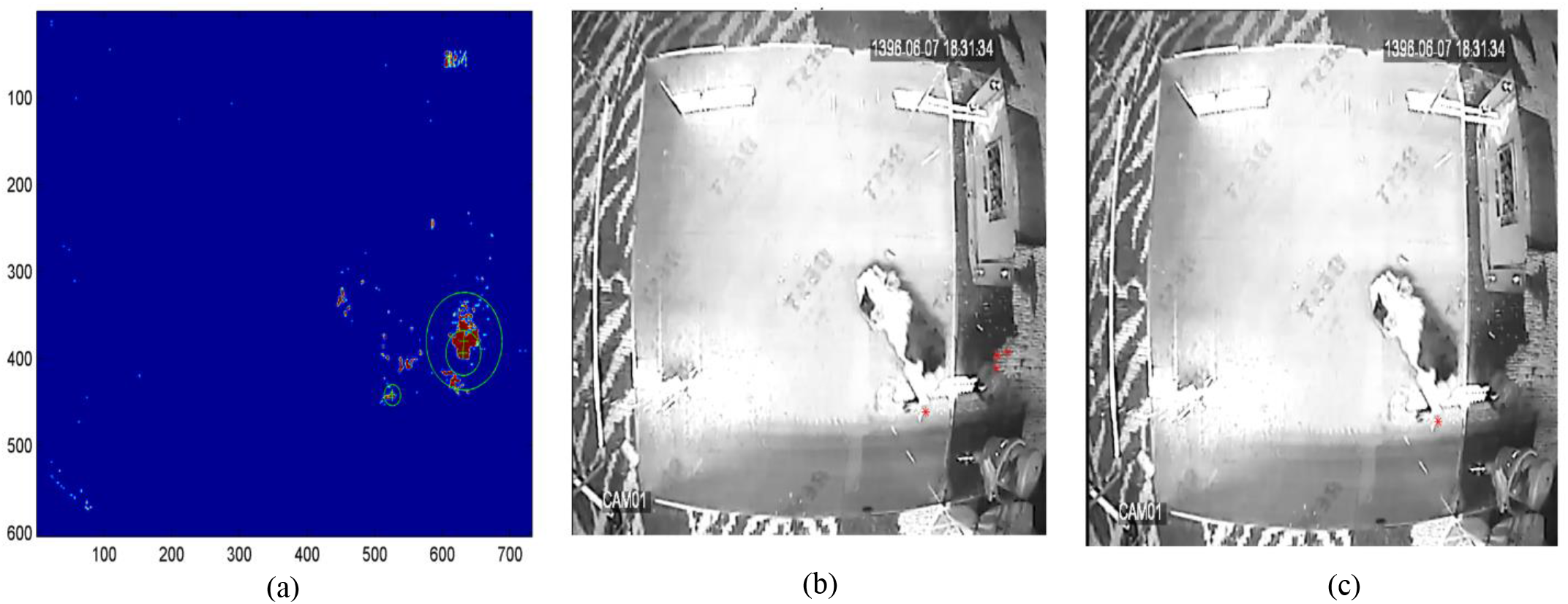
Result of (a) Wavelet transform and SURF feature points. (b) Location of SURF feature points in the mainframe. (c) Search for the animal only on the floor of the cage and tracked animal location

In order to plot animal movement diagrams, the time resolution of the tracking must first be determined. If the animal is moving fast, the position of the animal should be determined at small intervals, but if the animal is moving slowly, it will be sufficient to examine its behavior every second. Depending on the selected time accuracy, the number of points on the chart will also increase or decrease. Figure 7(a) shows the plot of animal movements for the time resolution of 2 seconds, Figure 7(b) shows for the resolution of 1 second and Figure 7(c) shows for the resolution of 0.5 seconds, all for the same video.

**Figure 7.**
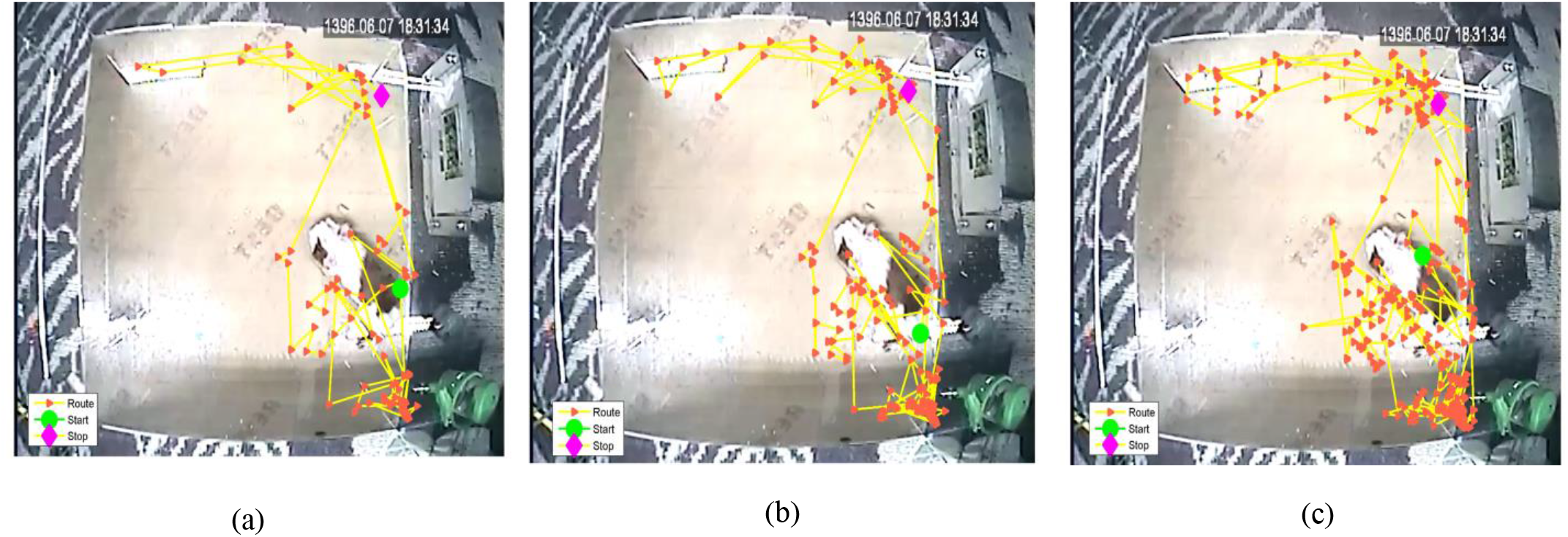
Drawing animal movement diagrams. (a) Once every 2 seconds. (b) Once every 1 second. (c) Once every 0.5 second.

### 3-2 auditory stimulation

To test the auditory stimulation system, a 5 V amplitude, 200 Hz sin wave stimulus is applied to the speaker in the cage for one minute, and the animal’s movements are monitored before, during and after the stimulation, as shown in Figure 8. Each episode has a frame rate of one frame per second, and each episode includes one minute of the video. According to the obtained results, it can be seen that the animal has a normal movement before the stimulation, its movement is minimized during the stimulation due to the animal’s fear, and after the stimulation, the animal moves more than the initial state.

**Figure 8.**
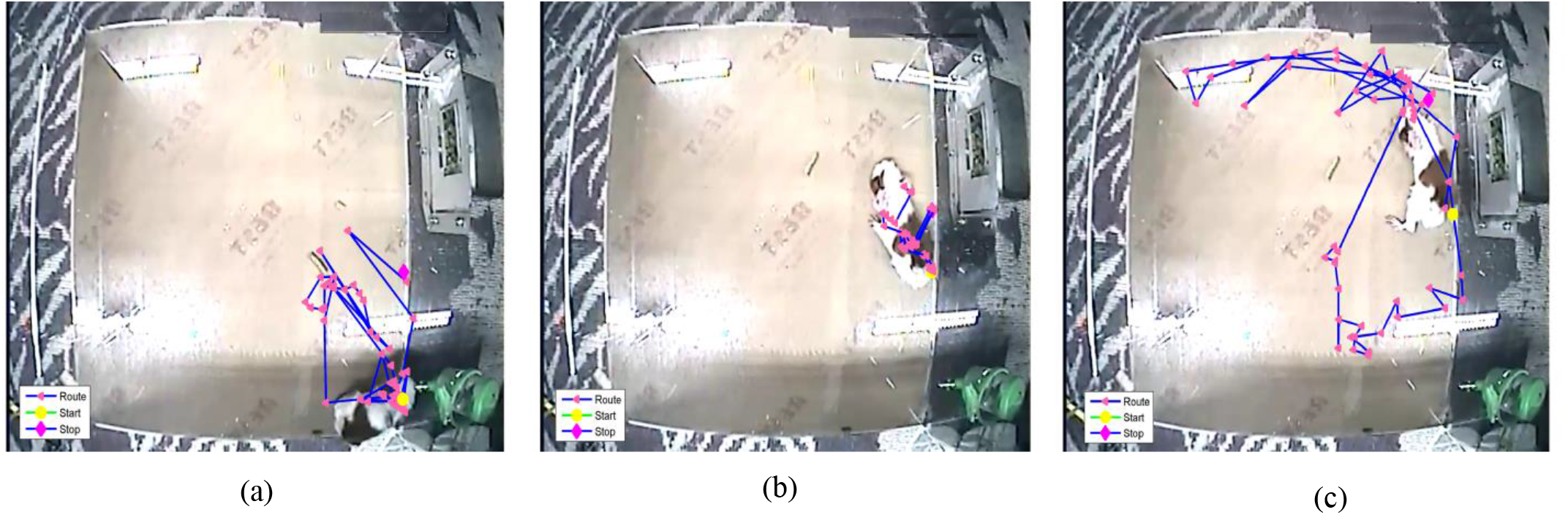
Investigation of animal movement behavior due to auditory stimulation. (a) Before the stimulation. (b) During the stimulation. (c) After the stimulation.

### 3-3 Cage temperature

To explore the temperature stability in the cage, we set the temperature to 25 degrees using the implemented hardware and measure the temperature in 24 hours. Figure 9 illustrates the temperature stability of this cage for 24 hours. Evidently, the installed hardware and the written code can keep the temperature of the cage almost constant for 24 hours and can minimize the temperature stress on the animal.

**Figure 9.**
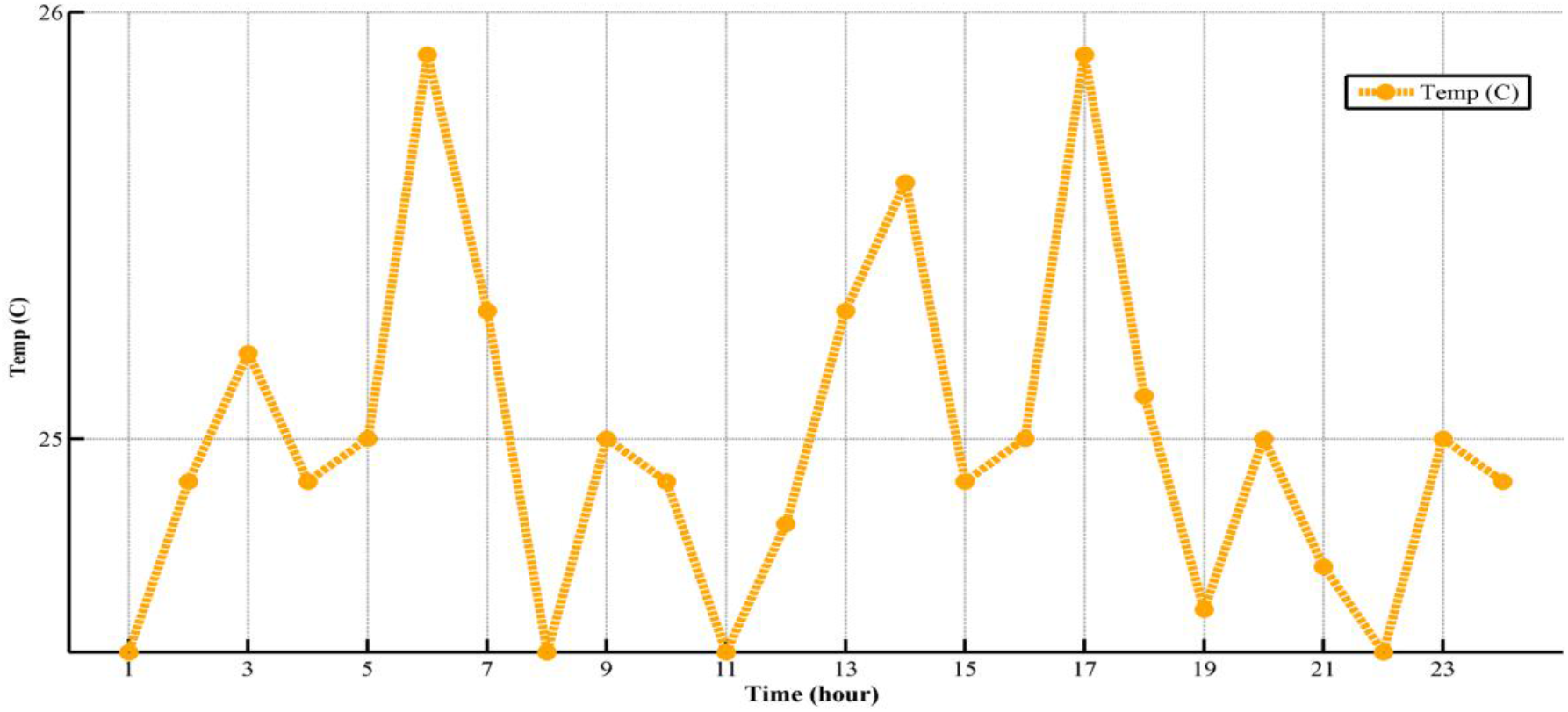
Temperature measured in the cage for 24 hours

## 4- DISCUSSION

In this section, we want to find the best pre-processing methods that can follow the animal’s movements properly. For this purpose, in the differential frame, we perform various pre-processing algorithms, then we extract the SURF descriptor and consequently the position of the animal. 5 number of one-minute video of the animal movements is randomly selected, then the position of the animal is determined once manually by the operator, and then by the proposed algorithm. One frame is processed every second of the animal’s movement (a total of 300 frames for 5 videos). The following performance parameters are defined to evaluate different methods:

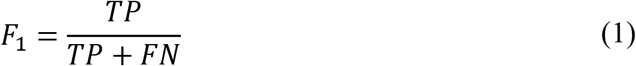

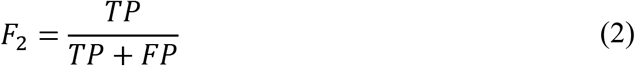

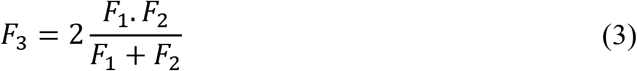

where TP (true positive) represents the number of animal positions that are correctly detected by the algorithm, FN (false negative) is the number of positions that are not detected, FP (false positive) is the number of situations that are incorrectly detected. Ideally, the values of F_1_, F_2_ and F_3_ should be calculated equal to 1. Table 1 shows the measured values presented in (1)–(3) and Figure 10 summarizes this table in the form of a graph.

**Figure 10.**
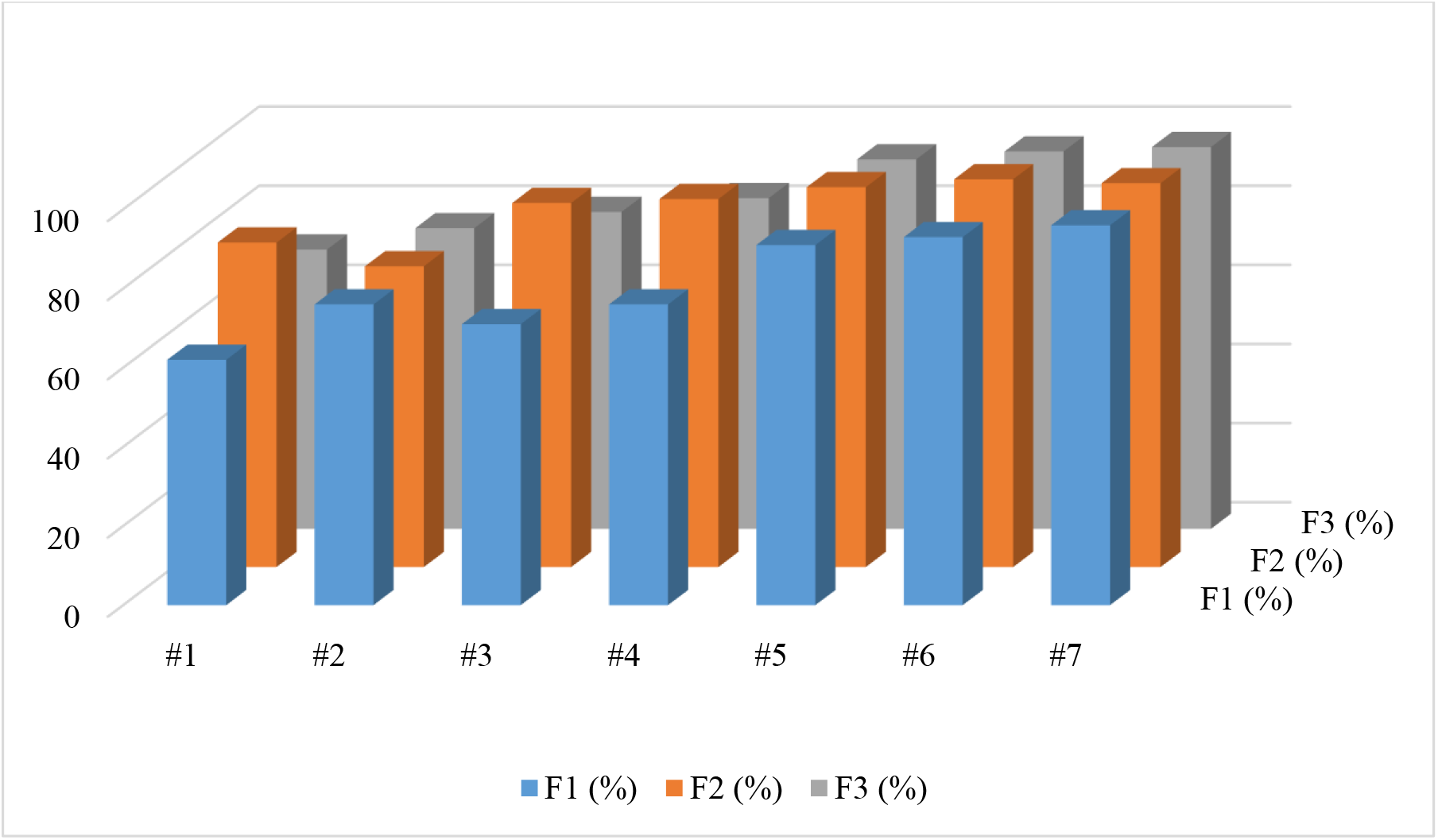
Results of different pre-processing methods and accuracy of animal position extraction.

**Table 1.** Different methods of video pre-processing and animal tracking performance

| Method | Type of pre-processing | TP | FN | FP | $F_1$ (%) | $F_2$ (%) | $F_3$(%) |
| --- | --- | --- | --- | --- | --- | --- | --- |
| #1 | Gamma mapping | 165 | 100 | 35 | 62 | 82 | 70.6 |
| #2 | Disk filter | 186 | 58 | 56 | 76 | 76 | 76 |
| #3 | Laplacian filter | 203 | 80 | 17 | 71 | 92 | 80.1 |
| #4 | Sobel filter | 218 | 68 | 14 | 76 | 93 | 83.6 |
| #5 | Fast Fourier Transform (FFT) | 265 | 25 | 10 | 91 | 96 | 93.4 |
| #6 | 2D Haar wavelet transform | 279 | 18 | 3 | 93 | 98 | 95.4 |
| #7 | 2D bior 3.7 wavelet transform | 285 | 9 | 6 | 96 | 97 | 96.5 |

According to the obtained results, the 2D bior 3.7 Wavelet transform method has the best accuracy in the position tracking. Analysis of the processed data also reveals that the main source of error in recognizing /not recognizing the position of the animal change in the brightness of the cage or the reflection of the animal on the wall of the cage.

### 4-1 Neuroscience Trials

The constructed cage was utilized for animal tests related to the treatment of Parkinson’s disease model in a rat with transcranial magnetic stimulation (TMS). Mobility, freezing, the traveled distance, and the grid test are inspected in that study, as shown in Figure 11. Details about the magnetic stimulation device can be found in [16] [17], and details of these tests can be found in [18] [19]. In that study, in addition to tracking the animal’s movements before and after the magnetic therapy, electroencephalogram (EEG) signals were also analysed, the details of the EEG processing algorithm can be found in [20].

**Figure 11.**
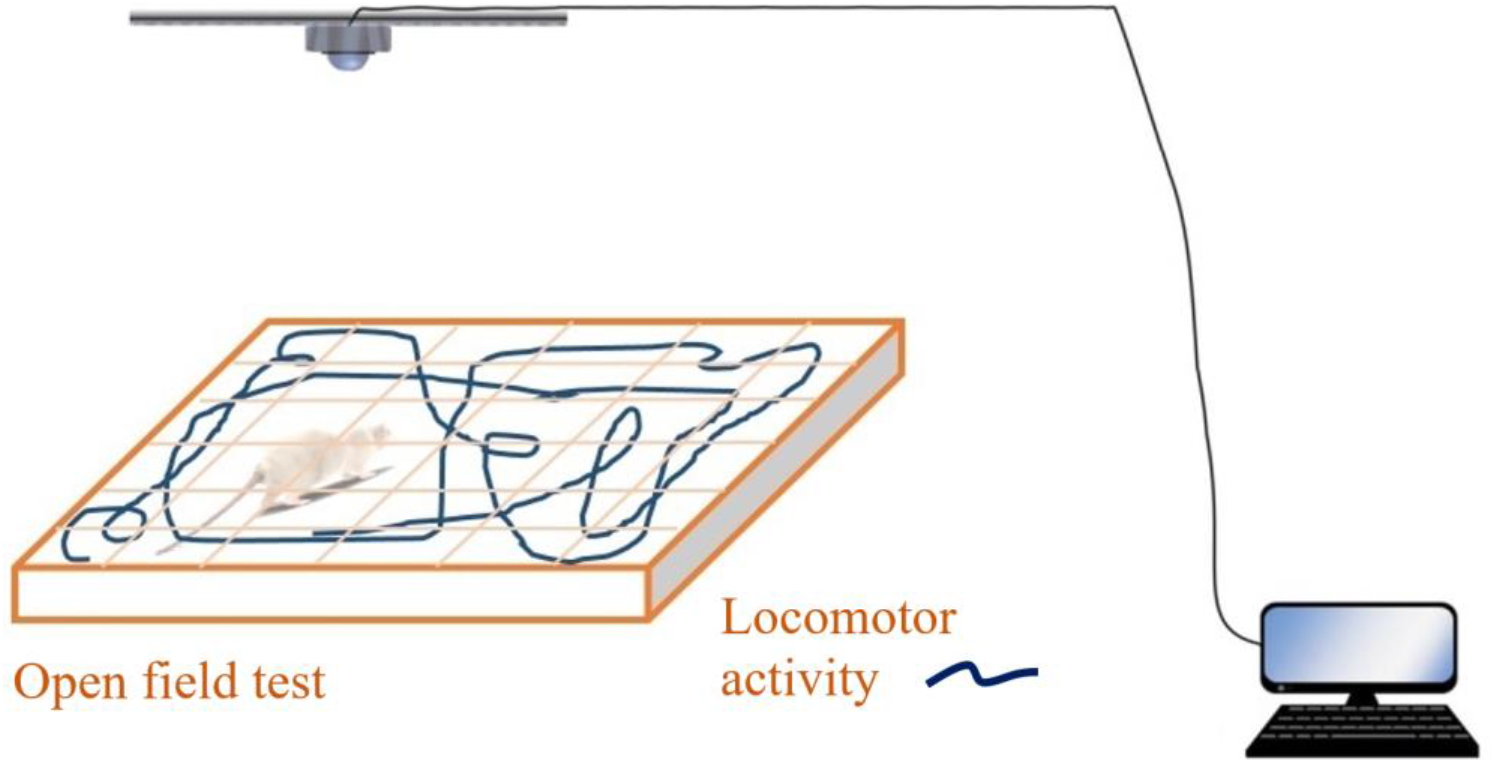
The animal movement tracking system for the tracking of hemi-Parkinsonian rat.

## 5- CONCLUSION

In this research, a novel cage based on image processing methods was designed and constructed to track the position of the laboratory animal in neuroscience trials. The main results of this study can be summarized as follows:

1. SURF feature points, along with discrete Wavelet transform, can be used as marker points in tracking laboratory animals. Therefore, the proposed algorithm in this research can be a good alternative to manual methods of tracking the animal.
2. The designed system and Fuzzy logic algorithm can control the thermal stability of the cage with an accuracy of ± 1 ° C.
3. The required lighting cycle can be adjusted with high accuracy.
4. The speaker installed in the cage can perform predefined auditory stimulations.

## Animal rights

All procedures performed in studies involving animals were in accordance with the ethical standards of the institutional and/or national research committee.

## References

[1] A. Gomez-Marin and et al., “Automated tracking of animal posture and movement during exploration and sensory orientation behaviors,” PloS one, vol. 7, no. 8, 2012.

[2] A. L. Samson and et al., “MouseMove: an open source program for semi-automated analysis of movement and cognitive testing in rodents,” Scientific reports, vol. 5, 2015.

[3] C. F. C. Junior and et al., “ETHOWATCHER: validation of a tool for behavioral and video-tracking analysis in laboratory animals,” Computers in Biology and Medicine, vol. 42, p. 257–264, 2012.

[4] L. P. J. J. Noldus and et al., “EthoVision: A versatile video tracking system for automation of behavioral experiments,” Behavior Research Methods, Instruments, & Computers, vol. 33, no. 3, pp. 398–414, 2001.

[5] M. Memarian Sorkhabi and M. Saadat Khajeh, “Classification of alveolar bone density using 3-D deep convolutional neural network in the cone-beam CT images: A 6-month clinical study,” Measurement, vol. 148, 2019/12/1.

[6] M. Memarian Sorkhabi and et al., “Physiological Artefacts and the Implications for Brain-Machine-Interface Design,” bioRxiv, 2020.

[7] M. Memarian Sorkhabi, “Computer Vision for Estimating Cooper Density by Optical Microscope Images,” American Journal of Computing Research Repository, vol. 2, no. 4, pp. 61–65, 2014.

[8] M. Memarian Sorkhabi, S. Bakhshi and S. Bashirian, “Computer Vision for Train Tracking System Using Discrete Wavelet Transform,” American Journal of Computing Research Repository, vol. 2, no. 3, pp. 53–57, 2014.

[9] “Chapter 35 - Ethical issues in animal biotechnology,” in Animal Biotechnology (Second Edition) Models in Discovery and Translation, Academic Press, 2020, pp. 709–729.

[10] C. Ware, “Chapter Eleven - Thinking With Visualizations,” in Information Visualization, Morgan Kaufmann, 2021, pp. 393–424.

[11] J. E. Schaik and et al., “Motion tracking in developmental research: Methods, considerations, and applications,” Progress in Brain Research, 2020.

[12] H. Morimitsu and et al., “Exploring structure for long-term tracking of multiple objects in sports videos,” Computer Vision and Image Understanding, vol. 159, pp. 89–104, 2017.

[13] E. Sangineto, “Pose and Expression Independent Facial Landmark Localization Using Dense-SURF and the Hausdorff Distance,” IEEE Transactions on Pattern Analysis and Machine Intelligence, vol. 35, no. 3, 2013.

[14] T. N. Shene and et al., “Real-Time SURF-Based Video Stabilization System for an FPGA-Driven Mobile Robot,” IEEE Transactions on Industrial Electronics, vol. 63, no. 8, 2016.

[15] M. León and et al., “How tall should a mink cage be? Using animals’ preferences for different ceiling heights to improve cage design,” Applied Animal Behaviour Science, pp. 24–34, 2017.

[16] M. Memarian Sorkhabi, J. Frounchi, P. Shahabi and H. Veladi, “Deep-Brain Transcranial Stimulation: A Novel Approach for High 3-D Resolution,” IEEE Access, vol. 5, pp. 3157–3171, 2017/2/23.

[17] M. Memarian Sorkhabi and et al., “Measurement of transcranial magnetic stimulation resolution in 3-D spaces,” Measurement, vol. 116, pp. 326–340, 2018.

[18] A. Mann and M.-F. Chesselet, “Chapter 8 - Techniques for Motor Assessment in Rodents,” in Movement Disorders (Second Edition), Academic press, 2015, pp. 139–157.

[19] M. Memarian Sorkhabi, J. Frounchi and P. Parehkari, “The Effect of Focused Transcranial Magnetic Stimulation on Behavioral Profiles and Motor Cortex Signals in the Animal Model of Parkinsonism,” SSRN3647545, 2020/7/9.

[20] M. Memarian Sorkhabi, “Emotion detection from EEG signals with continuous wavelet analyzing,” Am. J. Comput. Res. Repos, vol. 2, no. 4, pp. 66–70, 2014/12.

